# Changes in DNA methylation after trauma processing and meditation in a large group setting of 1.6 years duration (Timeless Wisdom Training)

**DOI:** 10.1101/2025.06.10.658799

**Authors:** Michael Grünwald, Sebastien De Landtsheer, Thomas Hübl, Thomas Sauter

**Affiliations:** Heubergstrasse 17, 83043 Bad Aibling, Germany; Department of Life Sciences and Medicine, University of Luxembourg, 6, avenue du Swing, L-4367 Belvaux, Luxembourg; Inner Science LLC, 2 Ranch Drive, Novato, CA 94945, USA

**Keywords:** Epigenetics, DNA methylation, trauma, individual process, group process, meditation, Timeless Wisdom Training

## Abstract

Psychological trauma is associated with significant alterations of biological functions and is correlated with epigenetic changes specifically of DNA methylation. Trauma therapy is aiming at relieving the impact of trauma and is in initial studies also correlated with changes in DNA methylation. In this study we explored the changes in whole blood DNA methylation of participants of a program focusing on individual, ancestral and collective trauma processing and meditation in a large group setting of 1.6 years duration. Based on accompanying questionnaires, training participants report slight improvements in anxiety, depression and overall life satisfaction and some mystical experiences. 3227 CpGs and 253 genes were found to be differentially methylated during the training. Although these genes are not involving any of the known trauma related genes and relevant gene ontology terms, they comprise a large number of genes involved in the nervous function, as well as in cellular and developmental functions, the immune system and metabolism. Also, epigenetic aging is predicted to slow down during training. In summary this pilot study yielded additional findings showcasing the potential correlation of trauma therapy and alterations of DNA methylation.

## Introduction

**Psychological trauma** can be defined as an emotional response to an event or an experience that is deeply distressing or disturbing. It can result from a wide range of situations, including accidents, natural disasters, violence, abuse, or significant loss. The impact of trauma can vary greatly among individuals, affecting their mental, emotional, and physical well-being (Merrill et al., 2025). Van der Kolk discusses trauma as a single event or an ongoing experience with impact on the animalistic part of the brain. Trauma thereby is not so much about the stories of what happened, but it is “the current imprint of that pain, horror, and fear living inside people”. He concluded that trauma may change our nervous system. An individual’s reactions to these events - the survival strategies - are based on sympathetic (fight, hyper-activated) or vagal reactions (flight, freeze, hypo-activated) (Van der Kolk, 2015).

**Trauma therapy** focuses on addressing both the psychological and physical effects of trauma, aiming to offer strategies that help alleviate symptoms like anxiety, depression, and PTSD (Porter, 2024). Among the non-pharmacological approaches Prolonged Exposure, Cognitive Behavioral Therapy and Cognitive Processing Therapy are recommended (Watkins et al., 2018), but a variety of other helpful approaches have been developed. Somatic Experiencing focuses on changing the sensations associated with traumatic experiences and has proven to be effective on a variety of symptoms (Kuhfuß et al., 2021; Payne et al., 2015) and can also successfully be applied in group psychotherapy settings (Taylor and Saint-Laurent, 2017). In therapy approaches based on the Polyvagal Theory the importance of co-regulation is emphasized, *i.e.* the importance of social connections and relationships in regulating the nervous system (Muller, 2010).

Traumatic experiences can result in **biological imprints in the body**. As summarized by Merrill et al, this can involve genetic and **epigenetic modifications**, changes in the brain structure, as well as alterations of hormone levels among others (Merrill et al., 2025). The epigenetic layer is therefore a key topic of research. Epigenetics encompasses heritable mechanisms to regulate gene expression that do not change the underlying genomic sequence (DNA). DNA methylation specifically is a heritable epigenetic mark which usually represses gene expression, plays an important role in development, aging and many diseases and has also been associated with early life stress and mental disorders. Trauma and stress related epigenetic modifications have been found for many genes, e.g. *FKBP5*, *MAOA*, *NR3C1*, *SLC6A4* and others have been associated with childhood trauma (Jiang et al., 2019) and post-traumatic stress disorder (Banerjee et al., 2017; Dell’Osso et al., 2023; Howie et al., 2019). Furthermore, changes of DNA methylation in association with trauma therapy have already been reported, *e.g.* Carvalho Silva et al. reported 110 differentially methylated regions related to effects of therapy in a longitudinal study (Carvalho Silva et al., 2024). Concordantly, epigenetic effects have also been reported for meditation practices as reviewed here (Kaliman, 2019). Sitting meditation was associated with DNA methylation changes on age-related CpG sites (Venditti et al., 2020). Despite these relevant findings, there is only limited understanding of trauma therapy approaches like Somatic Experiences on the DNA methylome.

The **Timeless Wisdom Training** (TWT) is a two-year program aimed at personal development and increased awareness, particularly in relation to individual, ancestral and collective traumatization. Trauma processing within this program is relationship-centered with the initial aim to develop safe relationships. Due to the increasing psychological safety in the group, the self-healing mechanism of the body, psyche and nervous system begin to become more active and initiate processing processes from the past. The TWT thereby integrates elements of Somatic Experiencing (Kuhfuß et al., 2021; Payne and Crane-Godreau, 2015), the NeuroAffective Relational Model (NARM) (Heller, 2012) and the Polyvagal Theory (Porges, 2023) and combines it with meditation and music with dance. Group counselling methods are used. A training group usually consists of 150 participants and meets in person around four times per year for four weeks altogether. Regular exchange in small groups and individual counselling as well as ongoing daily meditation practice complement the training.

In **this study** we evaluated the possible impact of trauma work within the TWT on the whole blood genome-wide DNA methylation of training participants versus an age and gender matched control group. This analysis was accompanied by a questionnaire-based assessment of general and mental health, positive and adverse childhood experiences, as well as mystical experiences.

## Results

To study the potential epigenetic and trauma related changes during the TWT, we recruited a group of healthy participants (training group, *n=57*), as well as a gender and age matched healthy control group (*n=57*).

### Training group reports more adverse and less positive childhood experiences

To assess an important starting point of the participants related to trauma, all participants were asked to fill in questionnaires (see methods) about their adverse (ACE, 10 questions) and positive childhood experiences (PCE, 14 questions). Interestingly, training participants reported clearly more ACEs indicating that they experienced on average 2.7 out of the listed ten ACEs compared to only reported 1.2 ACEs on average in the control group (see **Figure 1A**). The most reported ACEs thereby were insults by parents, missing support, divorce of the parents and mental illness in the household (not shown). Also, the number of participants without any ACEs was very low in the training group (6) compared to 31 in the control group. Accordingly, the experienced PCEs were lower in the training group compared to the control group with an average score of 2.3 across all 14 questions (rated from 1/Yes to 5/NO) against 1.8 in the control group (see **Figure 1B**). These findings could be rooted in a higher trauma load in the training group which could also enforce their motivation to participate in the training or a higher awareness of their childhood experiences.

**Figure 1:**
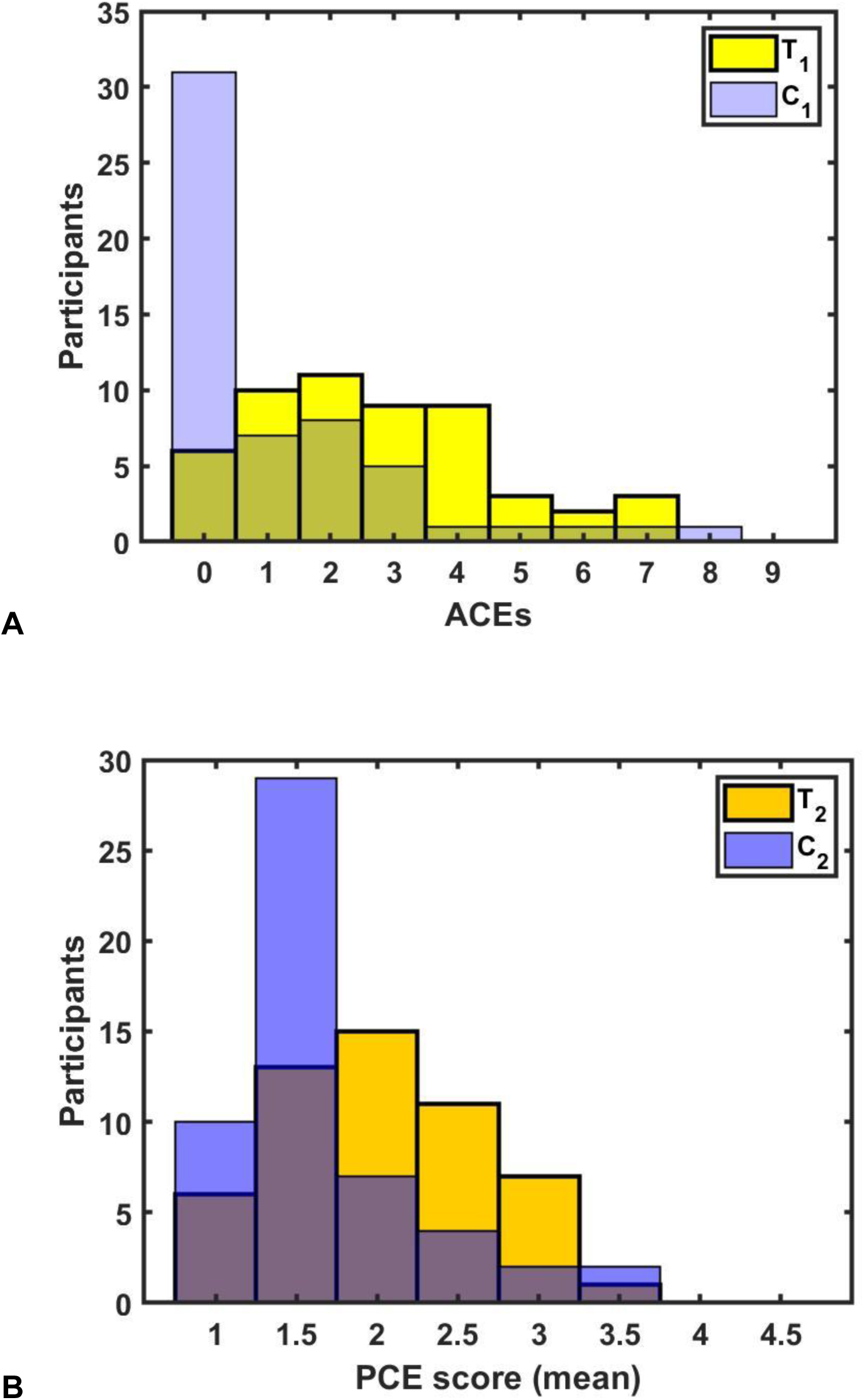
Training group reports more adverse and less positive childhood experiences. Training (*T*) and control group (*C*) participants filled the ACE-D questionnaire with 10 questions and the PCE questionnaire with 14 questions indicating if they made the listed adverse childhood experiences (YES/NO) or to what extent the listed positive childhood experiences (ranging from 1/Yes to 5/NO). **(A)** Number of participants (y-axis) of the training and control group who experienced a certain number (x-axis) of the listed 10 adverse childhood experiences. Higher values (x-axis) indicate more adverse childhood experiences. Data of time-point 1 are shown and are consistent with the data of time-point 2 (not shown). **(B)** Number of participants (y-axis) of the training and control group against the average score across all listed positive childhood experiences. Higher values (x-axis) indicate less positive childhood experiences. Data was obtained at time-point 2 only.

### Training group reports improvements in anxiety, depression and overall life satisfaction and some mystical experiences

For monitoring the changes in mental and general health, as well as in mystical experiences additional questionnaires were used at the beginning (time-point 1) and at the end of the training period (time-point 2). The data obtained with the PHQ-4 questionnaire (4 questions) revealed relatively low levels of anxiety and depression in both groups and further indicated a small but significant (*p=0.005*) decrease in anxiety and depression in the training group (see **Figure 2A**). Also, the life satisfaction reported within the EHIS questionnaire (106 questions) showed a small increase (not significant) in the training group while remaining constant at a high level in the control group (see **Figure 2B**). Remarkably, all the other questions of this general health questionnaire didn’t indicate any significant changes over the training period in the training versus the control group (not shown). Training participants reported high mystical experiences which further increased during the training period (*p=0.049*) in contrast to low and unchanged levels in the control group (see **Figure 2C**). Specifically, increased experiences of timelessness, of awesomeness, beyond description, and experiences without space boundaries and of an ultimate reality were mentioned by some training participants (not significant, not shown). Overall, these data show small but significant improvements in some mental and general health aspects and mystical experiences.

**Figure 2:**
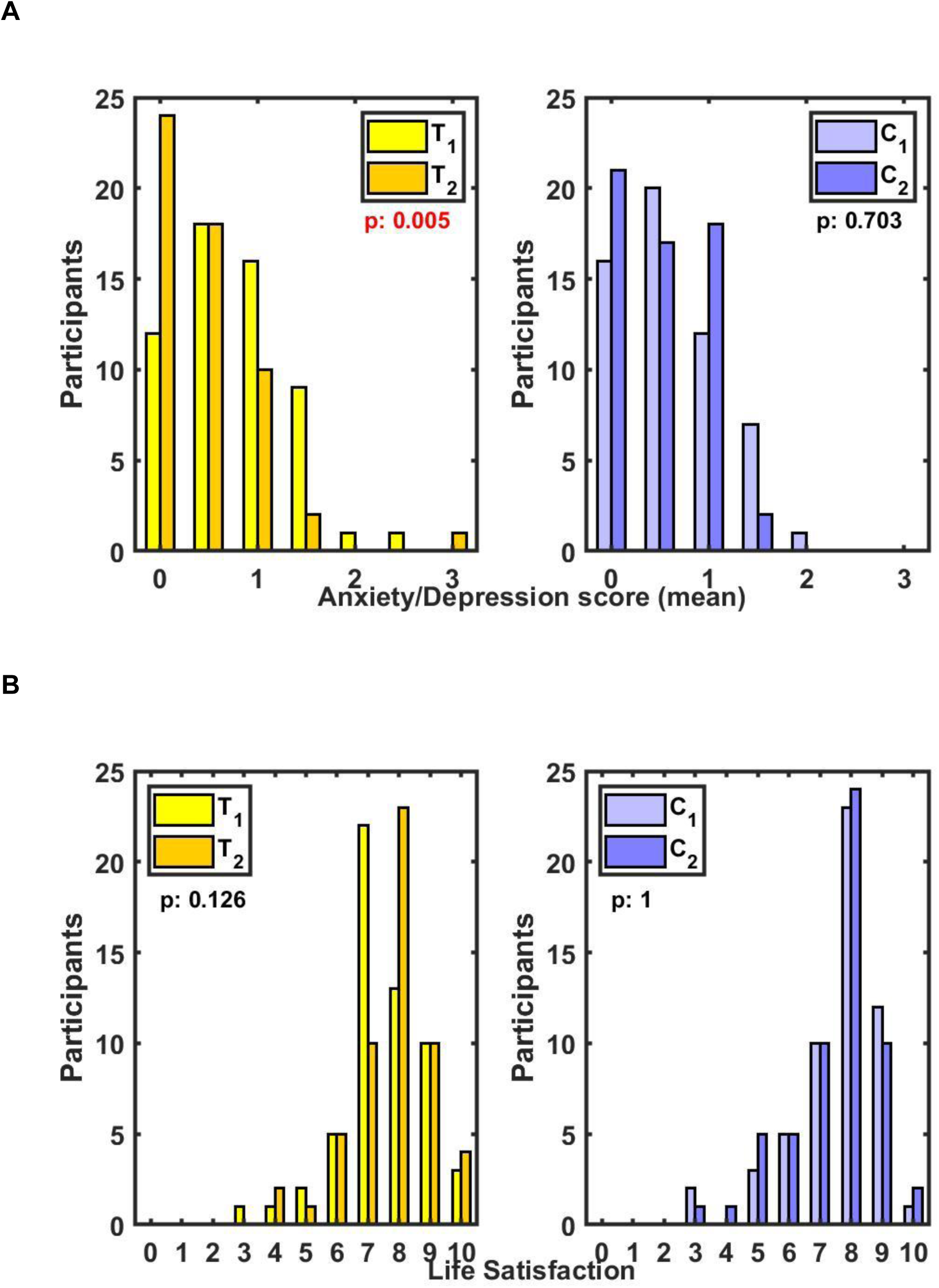
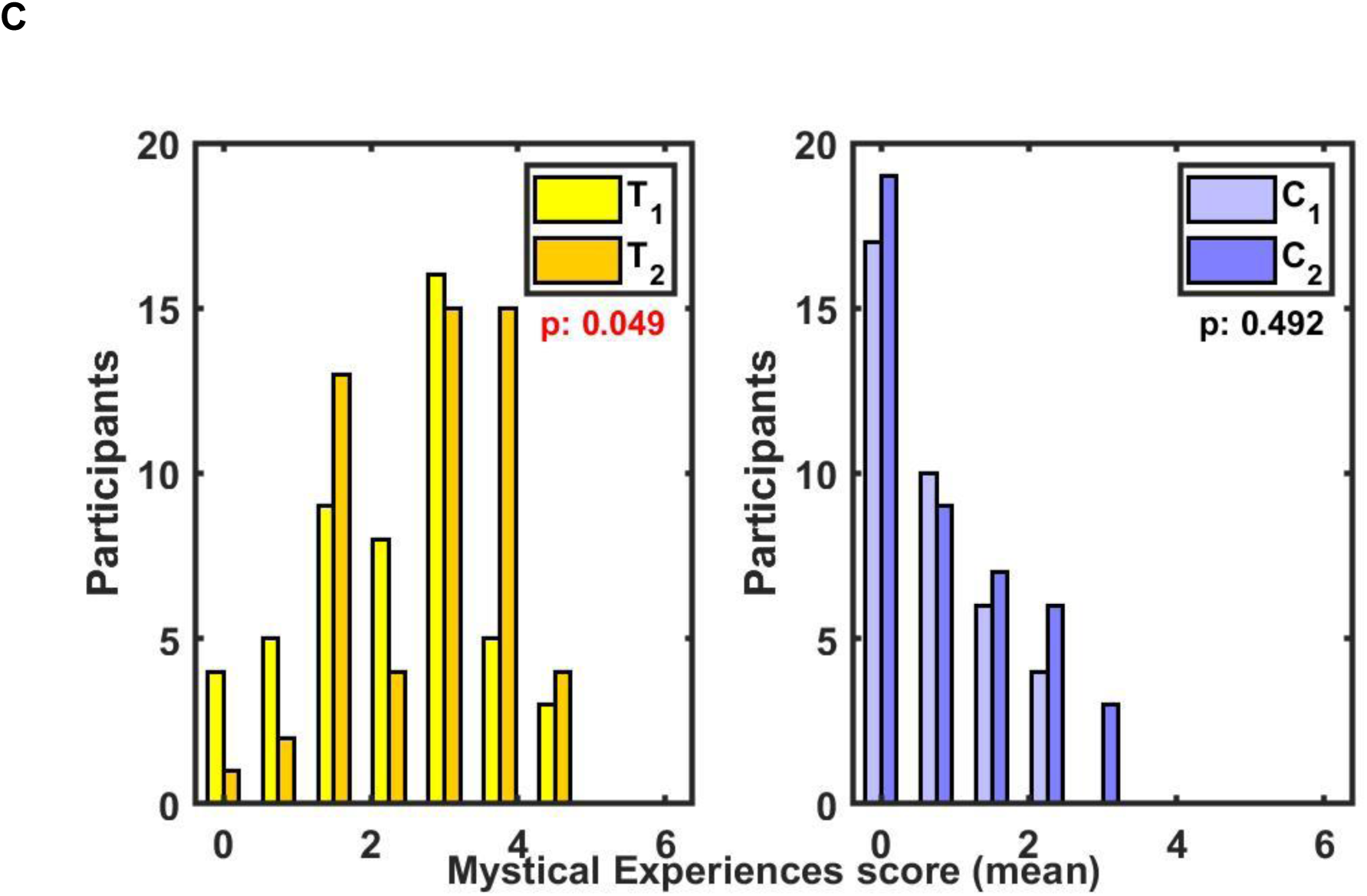
Training group reports improvements in anxiety and depression, as well as overall life satisfaction and some mystical experiences. Training (*T*) and control group (*C*) participants filled at time-point 1 and 2 the PHQ-4 questionnaire on anxiety and depression with 4 questions, EHIS questionnaire on general health with 106 questions and the MEQ30 questionnaire on mystical experiences. **(A)** Number of participants (y-axis) of the training (left panel) and control group (right panel) who indicated on time-point 1 and 2 a respective average score (x-axis) across all questions of the anxiety and depression questionnaire rated from 0/NO to 3/DAILY. Lower values (x-axis) indicate less experienced anxiety and depression. **(B)** Number of participants (y-axis) of the training (left panel) and control group (right panel) who indicated on time-point 1 and 2 in the EHIS questionnaire a respective life satisfaction (x-axis) rated from 1/LOW to 10/HIGH. Higher values (x-axis) indicate higher life satisfaction. **(C)** Number of participants (y-axis) of the training (left panel) and control group (right panel) who indicated on time-point 1 and 2 a respective average score (x-axis) across all questions of the mystical experiences questionnaire rated from 0/NO to 5/EXTREME. Higher values (x-axis) indicate more mystical experiences.

### 3227 CpGs and 253 genes found to be differentially methylated after training

To examine the potential effects of the TWT training on the epigenome (DNA methylation) of the participants, blood samples from the training and control group were collected at the beginning and at the end of the training period (*1.6 y*). DNA methylation levels of >850,000 CpG sites were determined with the Illumina EPIC2.0 BeadChip, followed by raw data processing including quality control, correction of a small batch effect in the control samples and differential methylation analysis (see methods). A rigorous selection method was applied focusing on consistent changes in the training group alone and in contrast comparing the training and control groups, but not in the control group (see methods). Thereby 3227 differentially methylated CpG sites could be determined (*adj.p-value<0.05*, see **Figure 3A** & **Supplementary Table 1**), showing increasing as well as decreasing methylation in the training group but not in the control group (see **Figure 3BC**). This high number of differentially CpG sites allowed determining 253 significant differentially methylated genes (see **Supplementary Table 2**). To elucidate the cellular functions covered by these genes, we first assembled from literature lists of genes of interest in trauma (105), childhood trauma (16) and resilience (35) research (see **Supplementary Table 3**). Surprisingly, no changes were observed in the methylation of neither this trauma and resilience related genes, nor in any of the CpG sites associated with these genes. This might be since all the participants were healthy and did not have any psychological diagnosis. Next, we performed general gene set enrichment analysis (see methods) and found some indications (not significant) for association of the significant genes with neural synaptic transmission (see **Figure 4AB**) and diseases linked to anxiety, tension and depression (see **Figure 4C**). To further consolidate this, we performed an extensive literature review which revealed a major implication of 80 of the differentially methylated genes in functions of the neuronal system (see **Supplementary Table 2**) and to a lesser extent in functions related to development, the immune system and metabolism, as well as others. In summary, 3227 CpG sites and 253 genes were found to be differentially methylated in the training group and - despite being widespread over the genome - with many of the genes being linked to functions of the neuronal system.

**Figure 3:**
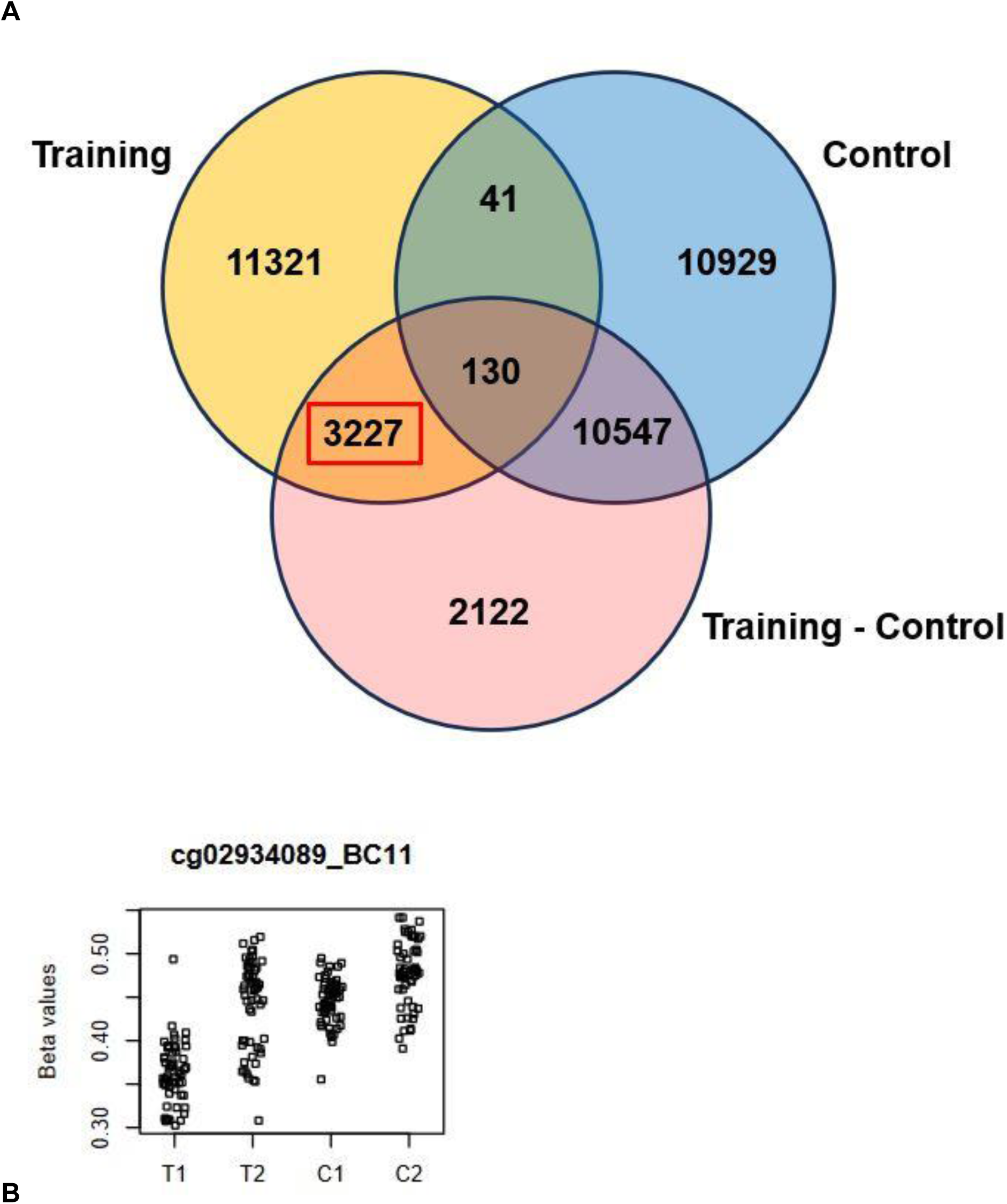
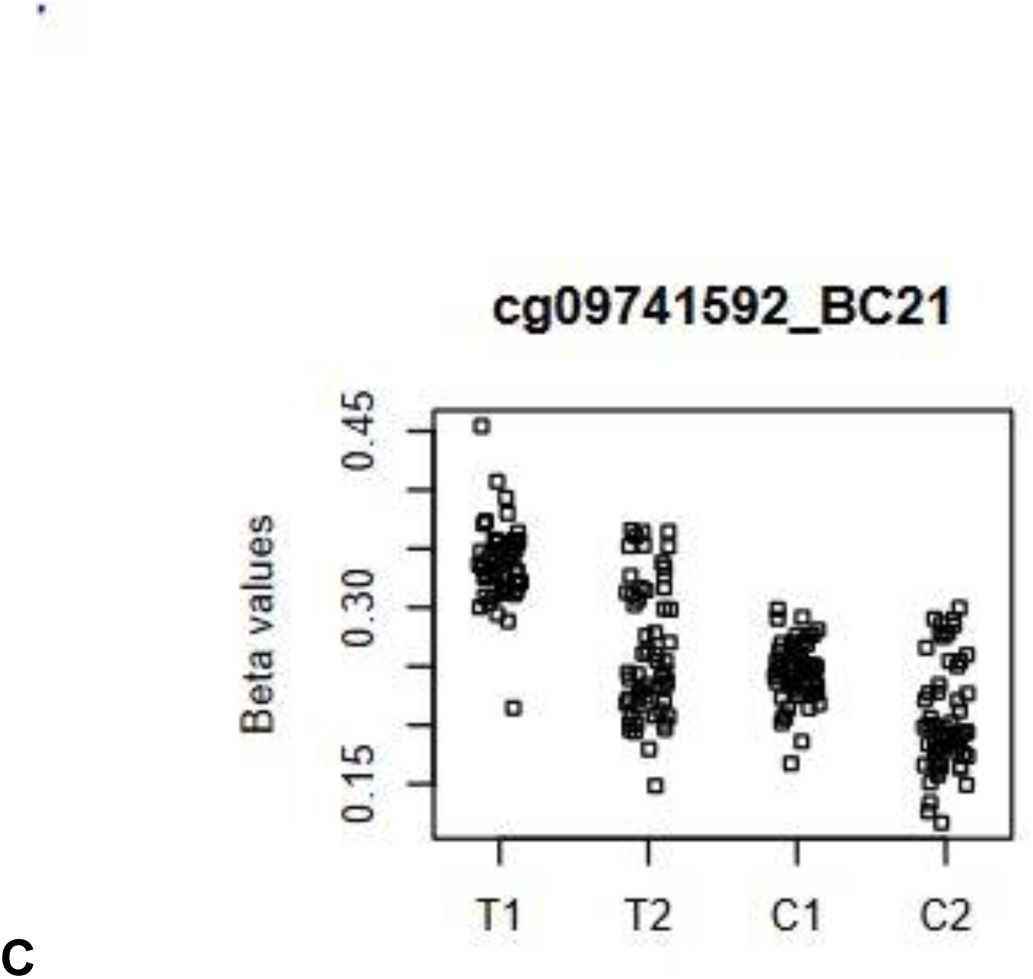
3227 CpGs found to be differentially methylated during training. Methylation at >850,000 CpG sites was determined with the EPIC2.0 BeadChip from blood samples of training (*T*) and control group (*C*) participants at time-point 1 and 2. **(A)** Number of unique and shared differentially methylated CpG sites between the two time-points (*FDR<0.05*) for the training group, the control group and the contrast between the training and control group (Training - Control, *cTC*, see methods). Red rectangle marks the called differentially methylated CpG sites found in the training group, in the Training - Control contrast, but not in the control group (see methods). **(B)** Example of a differentially methylated CpG site (beta-value, *n=57+57*) indicating a significant increase in the training group from time-point 1 (T_1_) to time-point 2 (T_2_). **(C)** Example of a differentially methylated CpG site (beta-value, *n=57+57*) indicating a significant decrease in the training group from time-point 1 (T_1_) to time-point 2 (T_2_).

**Figure 4:**
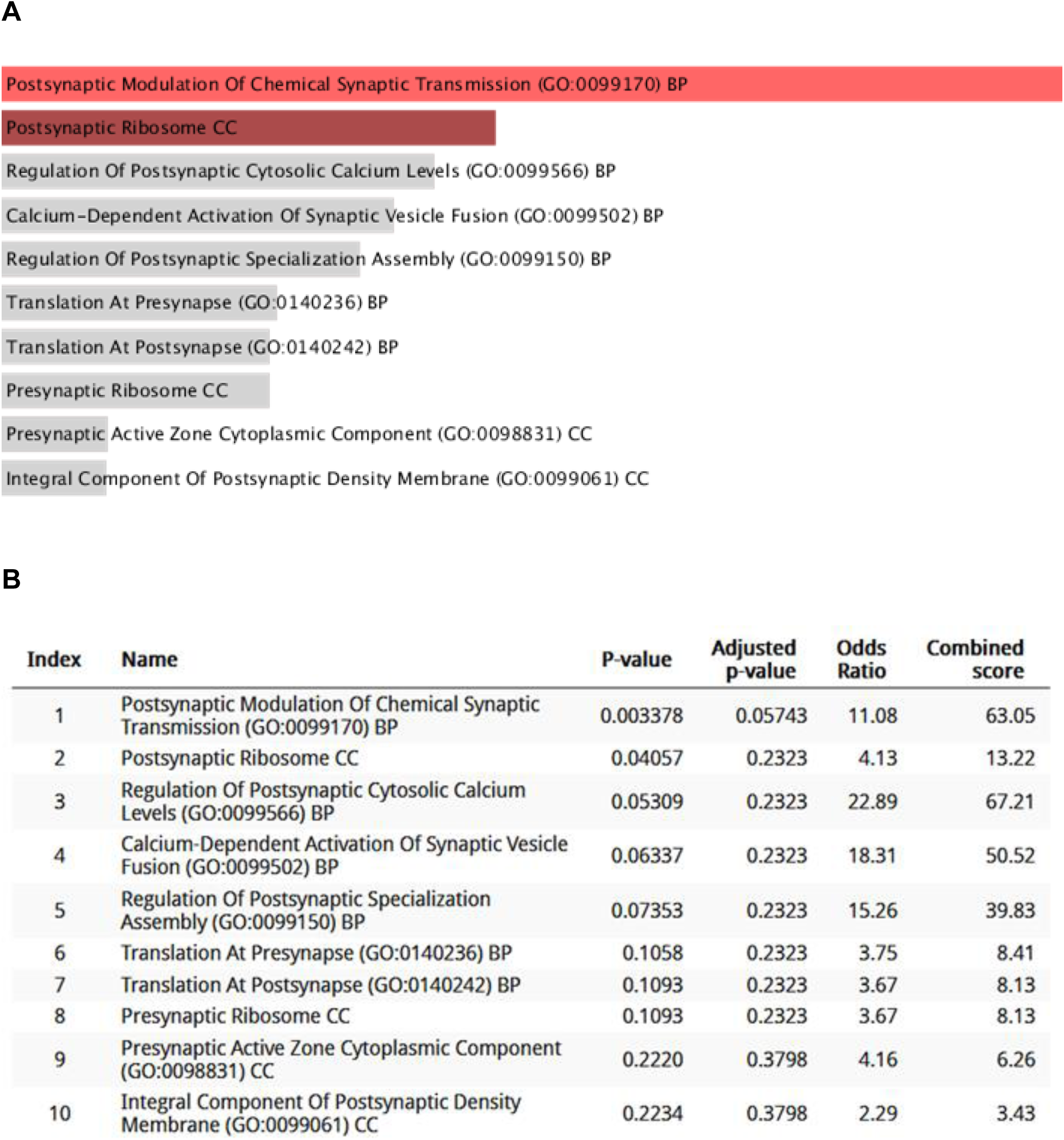
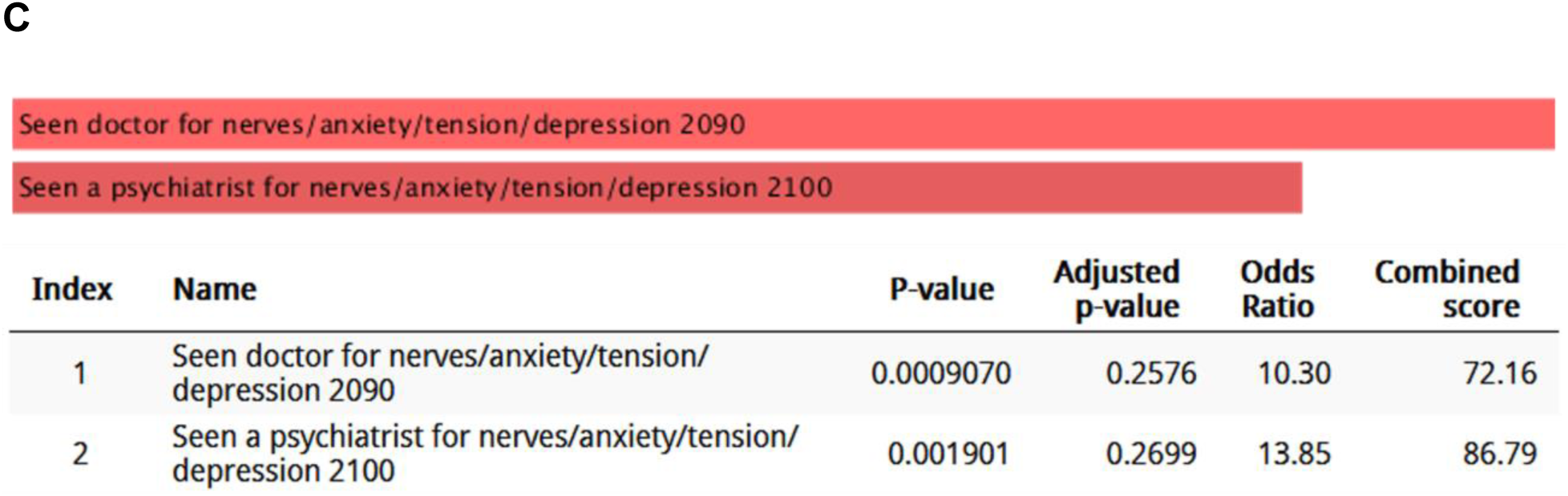
Pathway enrichment analysis of the differentially methylated genes shows potential (non-significant) links to neurons and mental diseases. Differentially methylated genes with annotated gene symbols (218) were subjected to pathway enrichment analysis with Enrichr. **(A, B)** Top 10 enriched GO terms in Enrichr - Ontologies - SynGO and their respective statistical values. Red bars indicate *p-value<0.05*. **(C)** Top 2 enriched GO terms in Enrichr - Diseases/Drugs - UK Biobank GWAS v1 and their respective statistical values. Red bars indicate *p-value<0.05*.

### Individual methylation changes found to be heterogeneous and linked to body mass index and potentially to training success

To further analyze the individual DNA methylation changes we first plotted per training participant the CpG sites, which changed at least 20% in their beta-values over the training period (see **Figure 5A** for the participant with most changes). This revealed heterogenous individual changes from 124 to 4359 CpG sites and based on this from 12 to 398 differentially methylated genes (see **Figure 5B**). When focusing on genes from this list which were shared by at least 25% of the training participants, 28 genes are found (see **Figure 5C**) which once again are enriched for genes of importance in neuronal cell types (see **Figure 5D**). Using the information of the differential methylation of these genes, clusters of participants could be identified with participants showing changes in many of these genes (responders, see **Figure 5C**, top and bottom clusters) and other participants with very few changes (non-responders, middle cluster). Responders thereby tend to have a slightly higher body-mass-index (BMI; mean=24.4 vs. 22.3 for non-responders) which might be linked to larger epigenetic drift. It has been shown that this slow but steady change of the epigenome is associated with higher BMI and accelerated aging (Foster et al., 2023; Hamilton et al., 2019; Issa, 2014). Interestingly, non-responders - besides often having a lower BMI - reported a stronger improvement in the anxiety and depression score (see **Figure 5C**). Besides giving interesting insights into the heterogeneity of individual DNA methylation changes over the training period, these analyses are statistically underpowered and would need more participants to reveal significant findings.

**Figure 5:**
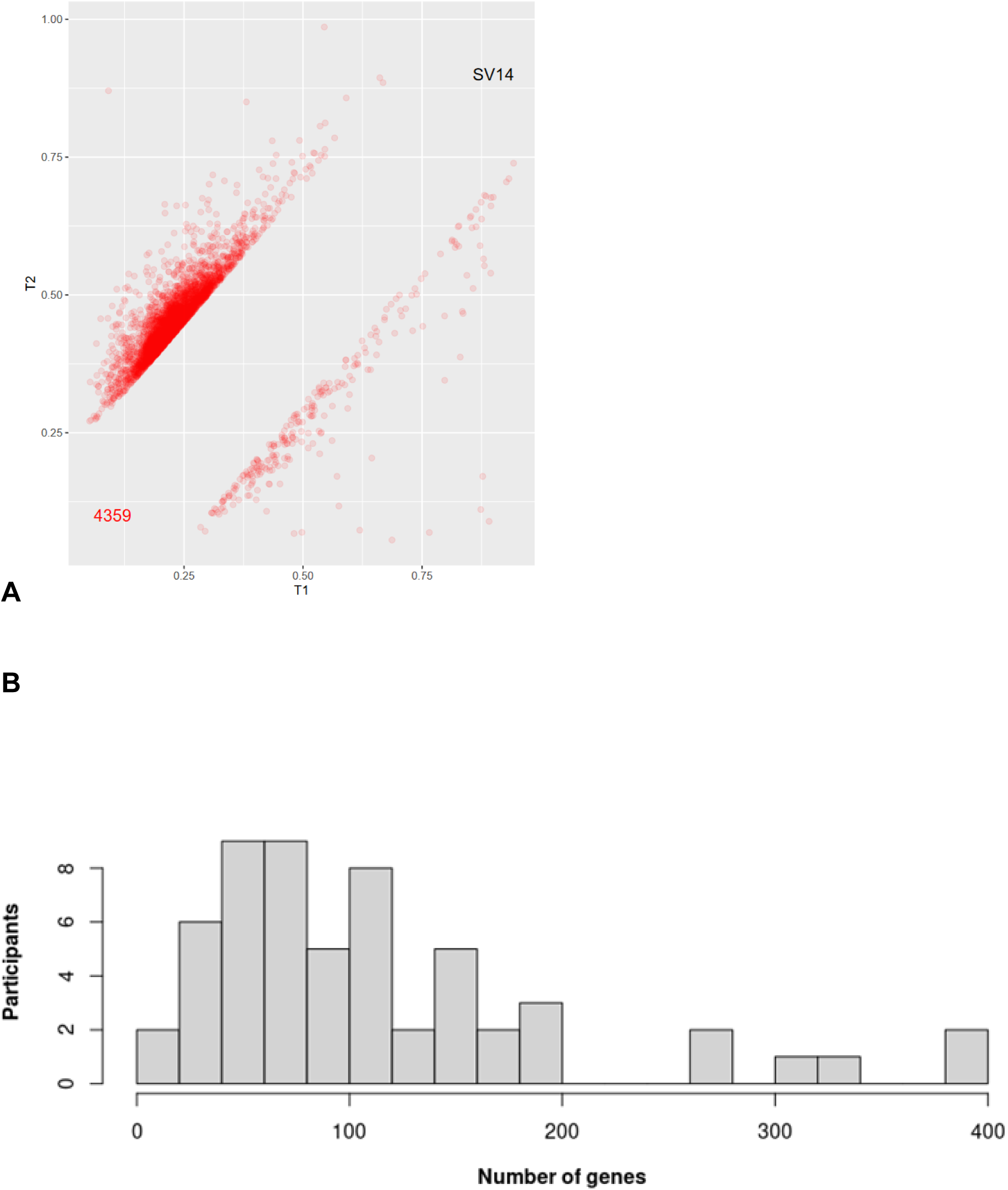
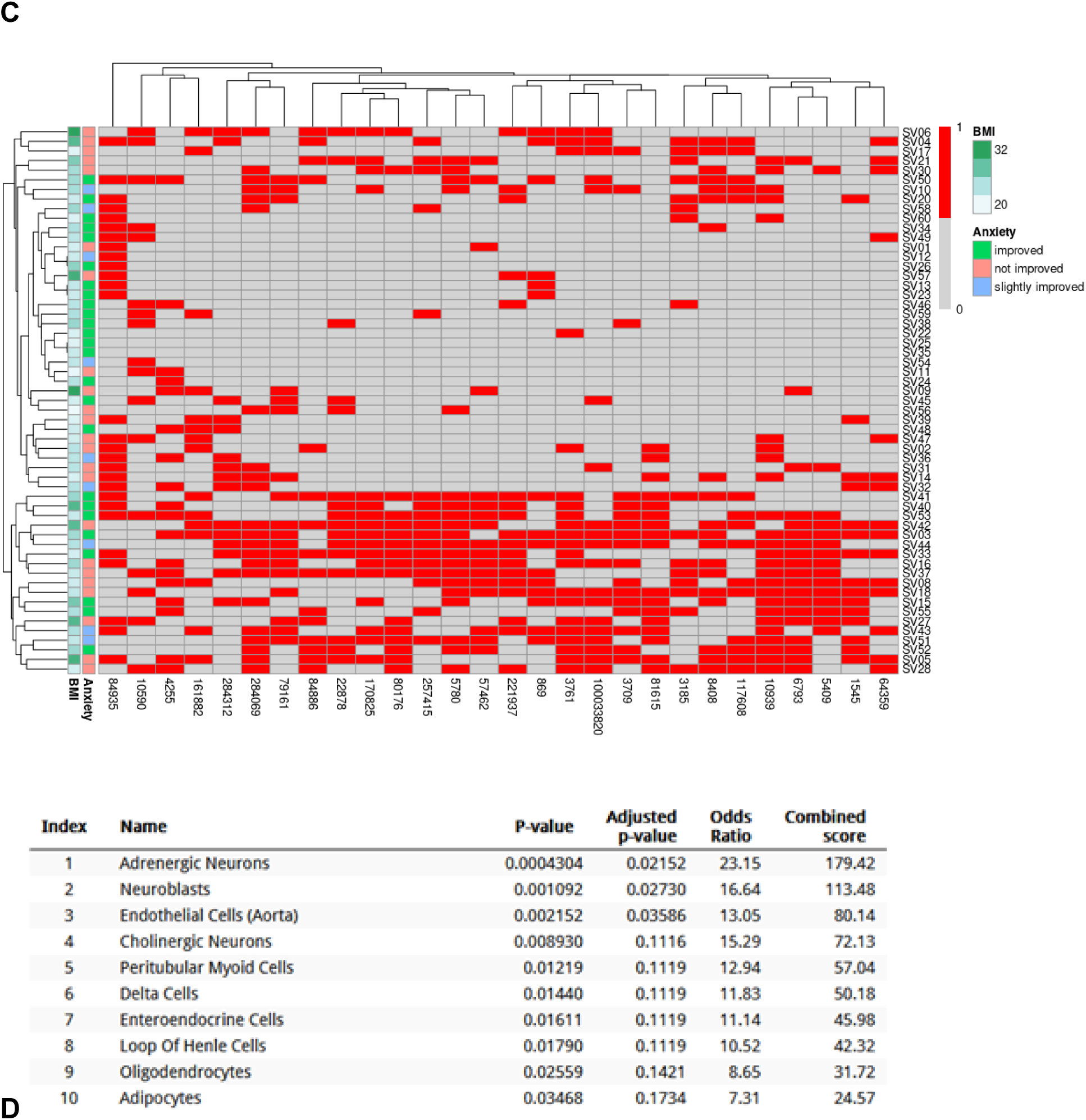
Individual methylation changes found to be heterogeneous and linked to body mass index and potentially to training success. Individual changes in DNA methylation of training participants were analyzed. **(A)** Scatterplot of beta-values of all CpG sites with at least 20% change between time-point 1 (T_1_) and time-point 2 (T_2_) for the training participant with most changes (“SV14”). Points in the upper-right corner indicate CpG sites with increased methylation over the training period. The red number indicates the number of (plotted) CpG sites with at least 20% change. **(B)** Number of participants (y-axis) of the training group against the number of differentially methylated per participant based on CpG sites with at least 20% change. Higher values (x-axis) indicate more individual methylation changes. **(C)** Clustering of individual differentially methylated genes (x-axis) found in at least 25% of the training participants (y-axis). Red indicates differential methylation in a respective participant. Annotation columns contain the body-mass-index (BMI) and the qualitative individual change in anxiety and depression over the training period (as determined by the PHQ-4 questionnaire average score, see Figure 2A) of the respective participant. **(D)** Statistical values of the top 10 enriched cell types in Enrichr - Cell Types - PanglaoDB Augmented 2021.

### Epigenetic aging is predicted to slow down during training

Finally, we examined the change of the predicted epigenetic age over the training period and across the training and study group. Epigenetic age acceleration is often associated with various diseases (Perna et al., 2016). We predicted the DNA methylation age with the Horvath epigenetic clock based on 353 CpG sites (Horvath, 2013). Despite a huge variability and uncertainty in the predicted values, a significant difference can be found between the training and control group. While the epigenetic age of the training participants increased on average by *1.3 y* over the 1.6-y training period, it increased by *3.3 y* in the control group (*p=0.005*, see **Figure 6A**). No differences were found between the responders and non-responders in terms of the shared differentially methylated genes in the individual analysis (see **Figure 5C** and **Figure 6B**). These initial results on a reduced age acceleration in the training group open interesting research directions but will need more data for a stronger validation.

**Figure 6:**
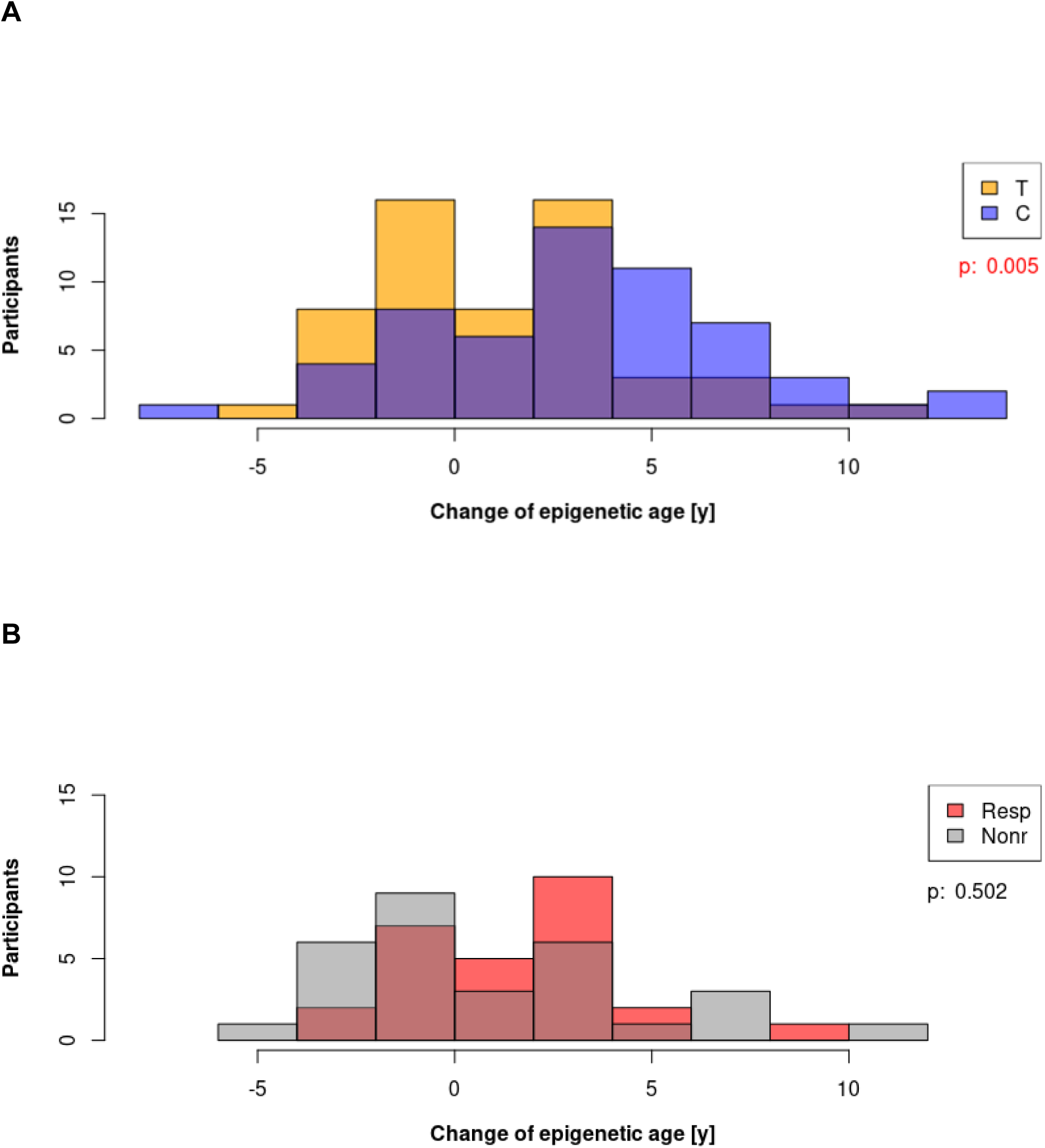
Epigenetic aging predicted to slow down during training. DNA methylation age was predicted for all participants at time-point 1 and 2. **(A)** Number of participants (y-axis) of the training (*T*) and control group (*C*) against their predicted change in epigenetic age [y] across the training period. **(B)** Number of training participants with individual changes in shared genes (Resp, responders, see Figure 5C, top and bottom clusters) and without such changes (Nonr, non-responders, see Figure 5C, middle cluster) against their predicted change in epigenetic age [y] across the training period.

## Discussion

Our study revealed some significant differences in DNA methylation and mental health of the TWT training participants across the training period compared to the age and gender matched control group. Based on the answers collected with five established questionnaires, it could be seen that the training participants report more adverse and less positive childhood experiences. It remains unclear whether this is indeed due to a higher burden during childhood or an increased awareness of such incidences. This, as well as the reported slight improvements in anxiety and depression during the training, would benefit from a more detailed examination with specifically elaborated questionnaires and at closer time intervals to give more conclusive insights.

Based on whole blood DNA methylation analysis, **3227 CpGs and 253 genes** were found to be differentially methylated in the participants of the TWT compared to the control group. The differentially methylated genes are mostly linked to the neuronal system, to development and regeneration, the immune system, metabolism and cellular and DNA associated processes (see **Supplementary Table 2**). Interestingly, it has already been shown elsewhere that blood transcriptome signatures are associated with molecular changes in the brain and clinical outcomes in Parkinson’s disease (Irmady et al., 2023). This mirroring of the brain transcriptome into the blood could be further analyzed in future work by performing a proteome analysis in the serum. In the following we discuss some of the overrepresented terms and relevant genes in the set of differentially expressed genes in more detail.

A large number (*n=80*) of the 253 differentially methylated genes are implicated in the **nervous system** and are related to its development, function and regeneration, *e.g. DCC* (DCC netrin 1 receptor) which is necessary for the development of the corpus callosum (Marsh et al., 2018) and the prefrontal cortex, including its dopamine innervation (Hoops and Flores, 2017), *SUN1* (Sad1 and UNC84 Domain Containing 1) associated with radial neuronal migration in the cerebral cortex (Zhang et al., 2009), *IPO7* (importin 7) implicated in peripheral nerve regeneration (Wen et al., 2022), *EGR4* (early growth response 4) which is found to be essential for the reconsolidation of previously established memories (Li et al., 2005) and *GJB6* (gap junction protein beta 6) which is associated with restoring auditory sensitivity via Connexin 26 (Zong et al., 2023).

Another large set of genes (*n=59*) is contributing to important **cellular and DNA associated processes,** *e.g. KCNQ1OT1* (KCNQ1 opposite strand/antisense transcript 1) is implicated in lineage specific gene silencing via DNA-methylation (Pandey et al., 2008), gene silencing (Kanduri et al., 2006) and epigenetic genomic imprinting (Soejima and Higashimoto, 2013)]. *ADAP1* (ArfGAP with dual PH domains 1) mediates cytoskeletal crosstalk via actin and microtubule cytoskeleton (Stricker and Reiser, 2014). *CMC2* (C-X9-C Motif Containing 2) is essential for cellular respiration as it is involved in the mitochondrial Cytochrome c oxidase biogenesis (Horn et al., 2010). *SMYD5* (SMYD family member 5) is known to have methyltransferase activity on histones (Rueda-Robles et al., 2021) and *SOD3* (Superoxide dismutase 3) is known to enhance overall cellular resilience (Kalmari and Colagar, 2024).

**Development and regeneration** associated genes (*n=37*) are *e.g. GPR160* (G protein-coupled receptor 160) regulating embryonic stem cell self-renewal (Fan et al., 2024), *KCNQ1OT1* (KCNQ1 opposite strand/antisense transcript 1) contributing to the epigenetic regulation of tissue development (Andresini et al., 2019; Azzi et al., 2014; Reeves, 1989), and *MMP17* (matrix metallopeptidase 17) involved embryogenesis, extracellular matrix remodeling, organogenesis, tissue regeneration, wound healing (Yip et al., 2019) and limb development (Blanco et al., 2017). *SMYD5* (SMYD family member 5) plays an important role in embryonic stem cell self-renewal and differentiation (Kidder et al., 2017), and *SRF* (serum response factor) contributes to the development of heart, vascular system, immune system, liver, neurons (Olson and Nordheim, 2010; Onuh and Qiu, 2021).

**Immune system** related genes (*n=27*) are found to be differentially methylated, e.g. *DDX1* (DEAD-box helicase 1) which is involved in antiviral immunity (Fullam and Schröder, 2013) and immunoglobulin class switching (Ribeiro de Almeida et al., 2018). *DEFB129* (defensin beta 129) contributes to anti-fungal (*C. albicans*) reactivity (Sakuma et al., 2022) and *MUC5B* (mucin 5B oligomeric mucus/gel-forming) to airway defense and mucociliary clearance (Roy et al., 2014) as key component of airway mucus (Rogliani et al., 2024).

**Metabolic processes** related genes (*n=25*) include *e.g. INSL5* (insulin like 5) which acts as an orexigenic hormone and modulates glucose homeostasis (Burnicka-Turek et al., 2012; Grosse et al., 2014; Hechter et al., 2022), *PON1* (paraoxonase 1) which effects HDL regulation (Mahrooz et al., 2019), HDL reverse transport and HDL antioxidant as well as in phosphotriester detoxification (Lewoń-Mrozek et al., 2024), and *SCAND1* (SCAN domain containing 1) which is involved in lipid metabolism (Babb and Bowen, 2003; West et al., 2013) and in the epigenetic programming of cardiovascular diseases in maternal diabetes (Higa et al., 2021). *SURF4* (surfeit 4) acts on α1-antitrypsin secretion, proinsulin secretion and VLDL secretion (Shen et al., 2023) and *C1QTNF9B* (C1q and TNF Related 9B) with the top statistical significance of *adj.p=2.1e-8* has a positive effect on fatty liver and lipid metabolism (Guan et al., 2022).

Besides these main groups also several other functions and relevant genes were found among the differentially methylated genes (see **Supplementary Table 2**). These include **heart** related genes (*n=24*), *e.g. CISD2* (CDGSH iron sulfur domain 2) which contributes to the delay of cardiac mitochondrial aging (Yeh et al., 2020), *TAS2R41* (taste 2 receptor member 41) and *IPO7* (importin 7) contributing to epicardiac adipose tissue (Wang et al., 2021) and heart regeneration (Wen et al., 2022). Among the **bone/cartilage/muscle** related genes (*n=21*) are *HOXD9* (homeobox D9) and *RPL3* (ribosomal protein L35) implicated in bone regeneration and chondrocyte senescence protection (J. Zhu et al., 2024), as well as *WNT5A* (Wnt family member 5A) and *HOXD9* (homeobox D9) involved in chondrogenesis (Inoue et al., 2024) and aging skeletal muscle methylation (Turner et al., 2020). **Blood/blood vessel** related genes (*n=20*) include *C1QTNF9B* (C1q And TNF Related 9B), acting coronary artery disease protective and endothelial anti-aging (Lee et al., 2021; Song et al., 2025; Z. Zhu et al., 2024), as well as *RGS18* (regulator of G protein signaling 18) involved in platelet reactivity (Ma et al., 2015). The **fertility** related genes *NOBOX* (NOBOX oogenesis homeobox) and *PDCL2* (phosducin like 2) contribute to oogenesis (Jagarlamudi and Rajkovic, 2012) and spermiogenesis (Li et al., 2022). *LINGO3* (leucine rich repeat and Ig domain containing 3), *PYDC1* (pyrin domain containing 1) and *NEAT1* (nuclear paraspeckle assembly transcript 1) are **stress** related genes which play a role in early childhood adversity epigenetics (Park et al., 2019), inflammation stress (Needham et al., 2015) and stress regulation in general (Kukharsky et al., 2020). And finally, some genes are linked to **longevity and aging** *e.g. CISD2* (CDGSH iron sulfur domain 2) which is essential for the delay of cardiac aging (Yeh et al., 2019) and prolongevity (Shen et al., 2021; Yeh et al., 2022) and *PDE4C* (phosphodiesterase 4C) in the context of epigenetic aging (Guvatova et al., 2023; Mongelli et al., 2023).

In summary, a high number of genes were found to be differentially methylated coherently across not only some individuals but in most of the training group participants with broad implications in the nervous system, sensory organs, development and regeneration, the immune system, cellular and metabolic processes, and aging. This list of genes might thus give an indication of the effects of individual, ancestral and collective trauma processing. Specifically, genes implicated in neuronal functions were found significantly differentially methylated. It is well known that neuronal plasticity can be guided by experience and be connected to early-life social adversities (Braun and Bogerts, 2001; Miskolczi et al., 2019) and that psychotherapy can contribute to rebuilding the brain (Malhotra and Sahoo, 2017). Interestingly, in the theory of human trauma and chronic stress as developed within Somatic Experiencing - one of the key therapy forms within this study - trauma is described as functional dysregulation of a core response network composed of subcortical autonomic, limbic, motor and arousal systems (Payne et al., 2015). Also, the polyvagal theory connects functions of the nervous system with trauma therapy (Manzotti et al., 2024). Such work may allow for halted evolutionary processes in childhood (Bock et al., 2014) to evolve and adapt to the present life situation (Klimecki et al., 2014, 2013; Singer et al., 2006), with present and ongoing regeneration of the brain (Malhotra and Sahoo, 2017) and positive effects on prefrontal neuronal circuits (Moss, 2025). Thus, it will in the long run be important to better understand (with additional experimental methods and questionnaires) if the related functions of the nervous system, immune system, regeneration and others are changed to a healthier level given the observed significant changes in DNA methylation of important genes.

## Methods

### Participants and blood sampling

Participants of the Timeless Wisdom Training (TWT) cohort 2022-23, as well as healthy individuals not involved in the training were invited to this study in early 2022. 57 individuals in the training group and 57 individuals in the control group completed the full study. The training (study) group *T* consisted of 16 men and 41 women in the age range between 26 and 69 years and with an average age of 45.5 years. The control group *C* was gender and age matched. We obtained written consent by all participants specifically for scientific purposes. The study was approved by the *Ethik-Kommission der Bayerischen Landesärztekammer*, Munich, Germany (Nr. 19082) and by the Ethics Review Panel of the University of Luxembourg (ERP 23-075 DNAmethy). Blood was drawn from all participants twice in the periods of March 2022 (time-point 1, *T*_1_ for training group) and April - June 2022 (time-point 1, *C*_1_ for control group) and of October 2023 (time-point 2, *T*_2_ for training group) and November 2023 - January 2024 (time-point 2, *C*_2_ for control group). The time between T1/C1 and T2/C2 was kept equal (1.6 y). Samples were stored at -20 degree Celsius and shipped together on dry ice to the Helmholtz Zentrum München, Neuherberg, Germany for DNA isolation and methylation measurement.

### DNA isolation and methylation array

DNA extraction from 3 ml of blood was performed using a magnetic bead-based method on a Chemagic 360 robot (Revvity, Lübeck, Germany). DNA was quantified using Qubit (Life Technologies, Darmstadt, Germany) and 750 ng of genomic DNA from each sample was bisulfite converted using the EZ-96 DNA Methylation Kit (Zymo Research, Orange, CA, USA). Subsequent methylation analysis of all samples was performed on an Illumina iScan platform (San Diego, CA, USA) using the Infinium MethylationEPIC v2.0 kit according to standard protocols provided by Illumina. Initial quality control procedures to run the assay and generate methylation data export files were performed using GenomeStudio version 2011.1 software with Module Methylation version 1.9.0.

### Processing of methylation data

Methylation data processing and analysis was performed using the R Bioconductor workflow package *methylationArrayAnalysis* (Maksimovic, J, 2025) mostly following the tutorial given here and using *minfi*, *missMethyl*, *DMRcate* and other methylation specific packages: https://www.bioconductor.org/packages/release/workflows/vignettes/methylationArrayAnalysis/inst/doc/methylationArrayAnalysis.html. In short, raw data are read in from IDAT files, as well as the annotation for the EPIC2.0 BeadChip (*IlluminaHumanMethylationEPICv2anno.20a1.hg38*). All samples passed the initial control by having a mean detection p-value of less than 0.05. Quantile normalization was then performed with *preprocessQuantile*, followed by removal of any probes that have failed in one or more samples, probes on the sex chromosomes, probes with SNPs at a CpG site as well as known cross reactive probes, resulting in 855,856 kept probes. Multidimensional scaling (MDS) was then performed with *plotMDS* followed by visual inspection for any batch effect. A small batch effect was detected for plate 3 mostly containing control samples and considered in the design matrix. Beta values were then calculated with *getBeta* and subjected to differential methylation analysis with *lmFit* with the design matrix containing treatment (*i.e.* training and control groups at time-point 1 vs. time-point 2), individual participant and plate for considering the batch effect. Three contrasts were built (i) cT = T_2_ - T_1_; (ii) cC = C_2_ - C_1_; (iii) cTC = (T_2_ - T_1_) - (C_2_ - C_1_), representing differentially methylated probes in the training group, in the control group and between the two groups. An *FDR* cutoff of 0.05 was applied, as well as the following selection strategy to avoid false positives due to the small batch effect in the control group samples: Probes significant in cTC and cT, but not in cC were called differentially methylated. Based on these, differentially methylated genes were determined by *getMappedEntrezIDs.* Gene ontology enrichment was performed with *gometh*, *gsameth* testing the *C2 curated gene sets* from MsigDB (Liberzon et al., 2015), as well as manually curated lists of genes related trauma (104), childhood trauma (16) and resilience (35) and *Enrichr* (Chen et al., 2013; Xie et al., 2021). For further analyzing individual participants, probes with a difference of at least 0.2 in the beta value were considered. Based on these, differentially methylated genes per participant were determined as mentioned above and clustered with *pheatmap* when found in at least 25% of the training participants. DNA methylation age was predicted with *predictAge* of the *SeSAMe* R package (Lee et al., 2024) and applying the Horvath epigenetic clock based on 353 CpG sites (Horvath, 2013).

### Processing of questionnaire data

To acquire accompanying data of the participants on their general and mental health, their adverse and positive childhood experiences as well as their mystical experiences, the following established questionnaires were used: (i) EHIS: General health (106 questions) (Hintzpeter et al., 2019); (ii) PHQ-4: Anxiety and depression (4 questions) (Kroenke et al., 2009); (iii) ACE-D: Adverse childhood experiences (10 questions) (Felitti et al., 1998); (iv) PCE: Positive childhood experiences (14 questions) (Kuhar and Zager Kocjan, 2021); (v) MEQ30: Mystical experiences (30 questions) (Maclean et al., 2012). Data was collected at the time of blood drawing (time-points 1 and 2; PCE only for time-point 2) with printed questionnaires and digitized manually. Data was plotted in MATLAB. Statistical tests were performed with *kstest2* (Two-sample Kolmogorov-Smirnov goodness-of-fit hypothesis test.). For visualization purposes, changes in anxiety and depression (PHQ-4) over the training period were discretized into improved (mean score decreased by at least 0.5), slightly improved (mean score decreased by 0.25) and not improved (means score constant or increased).

## Data availability

The data that support the findings of this study are not openly available due to reasons of sensitivity. Processed data generated during this study are available from the corresponding author on reasonable request. Supplementary Tables 1 and 3 are available from the corresponding author on request.

## Acknowledgment.

We would like to thank Dr. Peter Lichtner (Helmholtz Zentrum München, D-85764 Neuherberg, Germany) for performing the DNA methylation analysis with the EPIC2.0 chip, Thomas Hübl and Dr. Daphne Leinweber for contributing to the description of the TWT and Sabine Sauter and Anastasiia Avramenko for their help in digitizing the questionnaires.

