## Supplementary Table 2 for "Changes in DNA methylation after trauma processing and meditation in a large group setting of 1.6 years duration (Timeless Wisdom Training)"

| initial_alias | converted_alias | name | description | namespace |
|---|---|---|---|---|
| 100033820 | None | None | None | |
| 100129196 | ENSG00000186056 | MATN1-AS | MATN1 antisense RNA 1 [Source:H( | ENTREZGENE_ACC |
| 100130264 | ENSG00000179447 | SLC24A3-A: | SLC24A3 antisense RNA 1 [Source:H | ENTREZGENE_ACC |
| 100131303 | ENSG00000263201 | DPEP2NB | DPEP2 neighbor [Source:HGNC Sym | ENTREZGENE_ACC |
| 10022 | ENSG00000172410 | INSL5 | insulin like 5 [Source:HGNC Symbol; | ENTREZGENE_ACC |
| 100233209 | ENSG00000247774 | PCED1B-AS | PCED1B antisense RNA 1 [Source:H( | ENTREZGENE_ACC |
| 100288748 | ENSG00000261437 | LINC02894 | long intergenic non-protein coding | ENTREZGENE_ACC |
| 100302192 | ENSG00000221594 | MIR548F1 | microRNA 548f-1 [Source:HGNC Syr | ENTREZGENE_ACC |
| 100302218 | ENSG00000221520 | MIR1285-1 | microRNA 1285-1 [Source:HGNC Sy | ENTREZGENE_ACC |
| 100313884 | ENSG00000221616 | MIR548H4 | microRNA 548h-4 [Source:HGNC Sy | ENTREZGENE_ACC |
| 100313895 | ENSG00000221442 | MIR548F4 | microRNA 548f-4 [Source:HGNC Syr | ENTREZGENE_ACC |
| 100422913 | None | None | None | |
| 100505692 | None | None | None | |
| 100505811 | ENSG00000250427 | LINC02148 | long intergenic non-protein coding | ENTREZGENE_ACC |
| 100505817 | ENSG00000261780 | LINC02582 | long intergenic non-protein coding | ENTREZGENE_ACC |
| 100505893 | ENSG00000248228 | SLIT2-IT1 | SLIT2 intronic transcript 1 [Source:H | ENTREZGENE_ACC |
| 100506100 | ENSG00000223478 | ZDHHC12-[ | ZDHHC12 divergent transcript [Sour | ENTREZGENE_ACC |
| 100506107 | ENSG00000250968 | LINC02382 | long intergenic non-protein coding | ENTREZGENE_ACC |
| 100507657 | ENSG00000226741 | LINC02554 | long intergenic non-protein coding | ENTREZGENE_ACC |
| 100533467 | ENSG00000270181 | BIVM-ERCC | BIVM-ERCC5 readthrough [Source:H | ENTREZGENE_ACC |
| 100861506 | None | None | None | |
| 100861573 | ENSG00000224405 | LINC00572 | long intergenic non-protein coding | ENTREZGENE_ACC |
| 100874136 | ENSG00000225316 | TPTE2-AS1 | TPTE2 antisense RNA 1 [Source:HGI | ENTREZGENE_ACC |
| 100874205 | ENSG00000232684 | ATP11A-AS | ATP11A antisense RNA 1 [Source:H( | ENTREZGENE_ACC |
| 100996624 | ENSG00000229243 | LINC01981 | long intergenic non-protein coding | ENTREZGENE_ACC |
| 100996669 | None | None | None | |
| 101243545 | ENSG00000240567 | LINC02067 | long intergenic non-protein coding | ENTREZGENE_ACC |
| 101926948 | None | None | None | |
| 101926964 | ENSG00000231252 | None | novel transcript | ENTREZGENE_ACC |
| 101927003 | ENSG00000254266 | PKIA-AS1 | PKIA antisense RNA 1 [Source:HGN( | ENTREZGENE_ACC |
| 101927360 | ENSG00000234781 | LINC01103 | long intergenic non-protein coding | ENTREZGENE_ACC |
| 101927394 | ENSG00000227110 | LMCD1-AS: | LMCD1 antisense RNA 1 [Source:HG | ENTREZGENE_ACC |
| 101927406 | None | None | None | |
| 101927418 | None | None | None | |
| 101927592 | ENSG00000249345 | LINC02405 | long intergenic non-protein coding | ENTREZGENE_ACC |
| 101927641 | ENSG00000238217 | LINC01877 | long intergenic non-protein coding | ENTREZGENE_ACC |
| 101927657 | ENSG00000244791 | None | novel transcript | ENTREZGENE_ACC |
| 101927683 | ENSG00000236497 | LINC01744 | long intergenic non-protein coding | ENTREZGENE_ACC |
| 101927769 | ENSG00000226097 | None | novel transcript | ENTREZGENE_ACC |
| 101928015 | ENSG00000227260 | LINC01985 | long intergenic non-protein coding | ENTREZGENE_ACC |
| 101928062 | ENSG00000257815 | PRANCR | progenitor renewal associated non- | ENTREZGENE_ACC |
| 101928085 | ENSG00000240137 | ERICH6-AS: | ERICH6 antisense RNA 1 [Source:HG | ENTREZGENE_ACC |
| 101928136 | ENSG00000253686 | LINC01484 | long intergenic non-protein coding | ENTREZGENE_ACC |
| 101928163 | None | None | None | |
| 101928622 | ENSG00000250954 | None | novel transcript | ENTREZGENE_ACC |
| 101928767 | None | None | None | |
| 101928887 | ENSG00000224714 | LINC02671 | long intergenic non-protein coding | ENTREZGENE_ACC |
| 101929019 | ENSG00000251210 | LINC02270 | long intergenic non-protein coding | ENTREZGENE_ACC |
| 101929504 | None | None | None | |
| 101929559 | None | None | None | |
| 101929592 | ENSG00000229444 | ST3GAL3-A | ST3GAL3 antisense RNA 1 [Source:H | ENTREZGENE_ACC |
| 10224 | ENSG00000180855 | ZNF443 | zinc finger protein 443 [Source:HGN | ENTREZGENE_ACC |
| 10261 | ENSG00000140749 | IGSF6 | immunoglobulin superfamily memb | ENTREZGENE_ACC |
| 102723427 | ENSG00000225209 | None | novel transcript | ENTREZGENE_ACC |
| 102723692 | ENSG00000261448 | None | novel transcript, antisense to XYLT1 | ENTREZGENE_ACC |

| | | | | |
|---|---|---|---|---|
| 102725072 | ENSG00000293159 | POM121L1 | POM121 transmembrane nucleopo | ENTREZGENE_ACC |
| 10322 | ENSG00000135632 | SMYD5 | SMYD family member 5 [Source:HG | ENTREZGENE_ACC |
| 103689918 | None | None | None | |
| 103752589 | ENSG00000251179 | TMEM92-A | TMEM92 antisense RNA 1 [Source:I | ENTREZGENE_ACC |
| 10404 | ENSG00000104324 | CPQ | carboxypeptidase Q [Source:HGNC | ENTREZGENE_ACC |
| 10424 | ENSG00000164040 | PGRMC2 | progesterone receptor membrane ( | ENTREZGENE_ACC |
| 104355292 | ENSG00000235202 | LINC01525 | long intergenic non-protein coding | ENTREZGENE_ACC |
| 104413892 | ENSG00000231882 | F10-AS1 | F10 antisense RNA 1 [Source:HGNC | ENTREZGENE_ACC |
| 10527 | ENSG00000205339 | IPO7 | importin 7 [Source:HGNC Symbol;A | ENTREZGENE_ACC |
| 105369306 | ENSG00000233393 | None | novel transcript | ENTREZGENE_ACC |
| 105372999 | None | None | None | |
| 105376453 | ENSG00000224215 | C10orf67-A | C10orf67 antisense RNA 1 [Source:I | ENTREZGENE_ACC |
| 105377389 | ENSG00000224932 | LINC02262 | long intergenic non-protein coding | ENTREZGENE_ACC |
| 105377600 | None | None | None | |
| 105377622 | None | None | None | |
| 10567 | ENSG00000105404 | RABAC1 | Rab acceptor 1 [Source:HGNC Symb | ENTREZGENE_ACC |
| 105755953 | ENSG00000234752 | LINC02676 | long intergenic non-protein coding | ENTREZGENE_ACC |
| 10801 | ENSG00000184640 | SEPTIN9 | septin 9 [Source:HGNC Symbol;Acc: | ENTREZGENE_ACC |
| 10804 | ENSG00000121742 | GJB6 | gap junction protein beta 6 [Source | ENTREZGENE_ACC |
| 109616972 | ENSG00000201388 | SNORA68B | small nucleolar RNA, H/ACA box 68I | ENTREZGENE_ACC |
| 109617022 | ENSG00000206785 | SNORA15B | small nucleolar RNA, H/ACA box 15I | ENTREZGENE_ACC |
| 109623488 | None | None | None | |
| 10984 | ENSG00000269821 | KCNQ1OT1 | KCNQ1 opposite strand/antisense t | ENTREZGENE_ACC |
| 11033 | ENSG00000105963 | ADAP1 | ArfGAP with dual PH domains 1 [So | ENTREZGENE_ACC |
| 11170 | ENSG00000168309 | FAM107A | family with sequence similarity 107 | ENTREZGENE_ACC |
| 11193 | ENSG00000120688 | WBP4 | WW domain binding protein 4 [Sou | ENTREZGENE_ACC |
| 11224 | ENSG00000136942 | RPL35 | ribosomal protein L35 [Source:HGN | ENTREZGENE_ACC |
| 113157 | ENSG00000293352 | RPLP0P2 | ribosomal protein lateral stalk subu | ENTREZGENE_ACC |
| 114041 | None | None | None | |
| 114821 | ENSG00000232040 | SCAND3 | SCAN domain containing 3 [Source: | ENTREZGENE_ACC |
| 115004 | ENSG00000164430 | CGAS | cyclic GMP-AMP synthase [Source:I | ENTREZGENE_ACC |
| 117608 | ENSG00000178338 | ZNF354B | zinc finger protein 354B [Source:HG | ENTREZGENE_ACC |
| 120775 | ENSG00000178358 | OR2D3 | olfactory receptor family 2 subfamil | ENTREZGENE_ACC |
| 121355 | ENSG00000170627 | GTSF1 | gametocyte specific factor 1 [Sourc | ENTREZGENE_ACC |
| 124961 | ENSG00000180787 | ZFP3 | ZFP3 zinc finger protein [Source:HG | ENTREZGENE_ACC |
| 127396 | ENSG00000117010 | ZNF684 | zinc finger protein 684 [Source:HGN | ENTREZGENE_ACC |
| 127703 | ENSG00000142686 | C1orf216 | chromosome 1 open reading frame | ENTREZGENE_ACC |
| 128486 | ENSG00000197296 | FITM2 | fat storage inducing transmembran | ENTREZGENE_ACC |
| 1311 | ENSG00000105664 | COMP | cartilage oligomeric matrix protein | ENTREZGENE_ACC |
| 132200 | ENSG00000163632 | C3orf49 | chromosome 3 open reading frame | ENTREZGENE_ACC |
| 132954 | ENSG00000163440 | PDCL2 | phosducin like 2 [Source:HGNC Sym | ENTREZGENE_ACC |
| 1340 | ENSG00000126267 | COX6B1 | cytochrome c oxidase subunit 6B1 [ | ENTREZGENE_ACC |
| 135935 | ENSG00000106410 | NOBOX | NOBOX oogenesis homeobox [Sourc | ENTREZGENE_ACC |
| 140881 | ENSG00000125903 | DEFB129 | defensin beta 129 [Source:HGNC Sy | ENTREZGENE_ACC |
| 1428 | ENSG00000103316 | CRYM | crystallin mu [Source:HGNC Symbol | ENTREZGENE_ACC |
| 146562 | ENSG00000166246 | DNAAF8 | dynein axonemal assembly factor 8 | ENTREZGENE_ACC |
| 1588 | ENSG00000137869 | CYP19A1 | cytochrome P450 family 19 subfam | ENTREZGENE_ACC |
| 160728 | ENSG00000256870 | SLC5A8 | solute carrier family 5 member 8 [S | ENTREZGENE_ACC |
| 1630 | ENSG00000187323 | DCC | DCC netrin 1 receptor [Source:HGN | ENTREZGENE_ACC |
| 1653 | ENSG00000079785 | DDX1 | DEAD-box helicase 1 [Source:HGNC | ENTREZGENE_ACC |
| 170589 | ENSG00000149735 | GPHA2 | glycoprotein hormone subunit alph | ENTREZGENE_ACC |
| 191585 | ENSG00000280109 | PLAC4 | placenta enriched 4 [Source:HGNC S | ENTREZGENE_ACC |
| 195827 | ENSG00000158122 | PRXL2C | peroxiredoxin like 2C [Source:HGNC | ENTREZGENE_ACC |
| 1961 | ENSG00000135625 | EGR4 | early growth response 4 [Source:HG | ENTREZGENE_ACC |
| 196913 | ENSG00000214900 | LINC01588 | long intergenic non-protein coding | ENTREZGENE_ACC |
| 200523 | ENSG00000172073 | SPMIP9 | sperm microtubule inner protein 9 | ENTREZGENE_ACC |

| | | | | |
|---|---|---|---|---|
| 201158 | ENSG00000175106 | TVP23C | trans-golgi network vesicle protein | ENTREZGENE_ACC |
| 2041 | ENSG00000146904 | EPHA1 | EPH receptor A1 [Source:HGNC Sym | ENTREZGENE_ACC |
| 205327 | ENSG00000178074 | C2orf69 | chromosome 2 open reading frame | ENTREZGENE_ACC |
| 2110 | ENSG00000171503 | ETFDH | electron transfer flavoprotein dehy | ENTREZGENE_ACC |
| 2197 | ENSG00000149806 | FAU | FAU ubiquitin like and ribosomal pr | ENTREZGENE_ACC |
| 219844 | ENSG00000198331 | HYLS1 | HYLS1 centriolar and ciliogenesis as | ENTREZGENE_ACC |
| 220081 | ENSG00000165837 | ERICH6B | glutamate rich 6B [Source:HGNC Sy | ENTREZGENE_ACC |
| 220158 | ENSG00000263417 | GTSCR1 | Gilles de la Tourette syndrome chro | ENTREZGENE_ACC |
| 221946 | None | None | None | |
| 22986 | ENSG00000156395 | SORCS3 | sortilin related VPS10 domain conta | ENTREZGENE_ACC |
| 23141 | ENSG00000176915 | ANKLE2 | ankyrin repeat and LEM domain co | ENTREZGENE_ACC |
| 23172 | ENSG00000165660 | ABRAXAS2 | abraxas 2, BRISC complex subunit [ | ENTREZGENE_ACC |
| 23732 | ENSG00000260230 | FRRS1L | ferric chelate reductase 1 like [Sour | ENTREZGENE_ACC |
| 2519 | ENSG00000001036 | FUCA2 | alpha-L-fucosidase 2 [Source:HGNC | ENTREZGENE_ACC |
| 257629 | ENSG00000175311 | ANKS4B | ankyrin repeat and sterile alpha mo | ENTREZGENE_ACC |
| 25796 | ENSG00000130313 | PGLS | 6-phosphogluconolactonase [Sourc | ENTREZGENE_ACC |
| 259287 | ENSG00000221855 | TAS2R41 | taste 2 receptor member 41 [Sourc | ENTREZGENE_ACC |
| 260434 | ENSG00000169900 | PYDC1 | pyrin domain containing 1 [Source:I | ENTREZGENE_ACC |
| 262 | ENSG00000123505 | AMD1 | adenosylmethionine decarboxylase | ENTREZGENE_ACC |
| 26254 | ENSG00000188770 | OPTC | opticin [Source:HGNC Symbol;Acc:H | ENTREZGENE_ACC |
| 266971 | None | None | None | |
| 26996 | ENSG00000173890 | GPR160 | G protein-coupled receptor 160 [So | ENTREZGENE_ACC |
| 27134 | ENSG00000105289 | TJP3 | tight junction protein 3 [Source:HG | ENTREZGENE_ACC |
| 27288 | ENSG00000170748 | RBMXL2 | RBMX like 2 [Source:HGNC Symbol; | ENTREZGENE_ACC |
| 27442 | ENSG00000241832 | CECR3 | cat eye syndrome chromosome reg | ENTREZGENE_ACC |
| 283131 | ENSG00000245532 | NEAT1 | nuclear paraspeckle assembly trans | ENTREZGENE_ACC |
| 283150 | ENSG00000176302 | FOXR1 | forkhead box R1 [Source:HGNC Sym | ENTREZGENE_ACC |
| 284454 | None | None | None | |
| 284661 | ENSG00000235054 | LINC01777 | long intergenic non-protein coding | ENTREZGENE_ACC |
| 284825 | ENSG00000232079 | LINC01697 | long intergenic non-protein coding | ENTREZGENE_ACC |
| 286749 | ENSG00000068781 | STON1-GTF | STON1-GTF2A1L readthrough [Sour | ENTREZGENE_ACC |
| 29952 | ENSG00000176978 | DPP7 | dipeptidyl peptidase 7 [Source:HGN | ENTREZGENE_ACC |
| 309 | ENSG00000197043 | ANXA6 | annexin A6 [Source:HGNC Symbol;A | ENTREZGENE_ACC |
| 3217 | ENSG00000260027 | HOXB7 | homeobox B7 [Source:HGNC Symbo | ENTREZGENE_ACC |
| 3235 | ENSG00000128709 | HOXD9 | homeobox D9 [Source:HGNC Symbo | ENTREZGENE_ACC |
| 3293 | ENSG00000130948 | HSD17B3 | hydroxysteroid 17-beta dehydroger | ENTREZGENE_ACC |
| 3315 | ENSG00000106211 | HSPB1 | heat shock protein family B (small) | ENTREZGENE_ACC |
| 338653 | ENSG00000229414 | KCNQ1-AS1 | KCNQ1 antisense RNA 1 [Source:HG | ENTREZGENE_ACC |
| 339978 | ENSG00000250456 | LINC02260 | long intergenic non-protein coding | ENTREZGENE_ACC |
| 340508 | ENSG00000293209 | GAS2L1P2 | growth arrest specific 2 like 1 pseu | ENTREZGENE_ACC |
| 3440 | ENSG00000188379 | IFNA2 | interferon alpha 2 [Source:HGNC Sy | ENTREZGENE_ACC |
| 345274 | ENSG00000145283 | SLC10A6 | solute carrier family 10 member 6 [ | ENTREZGENE_ACC |
| 347240 | ENSG00000186638 | KIF24 | kinesin family member 24 [Source:H | ENTREZGENE_ACC |
| 3761 | ENSG00000168135 | KCNJ4 | potassium inwardly rectifying chan | ENTREZGENE_ACC |
| 388830 | ENSG00000237664 | LINC00316 | long intergenic non-protein coding | ENTREZGENE_ACC |
| 389043 | None | None | None | |
| 390651 | ENSG00000291191 | OR4F13P | olfactory receptor family 4 subfami | ENTREZGENE_ACC |
| 3984 | ENSG00000106683 | LIMK1 | LIM domain kinase 1 [Source:HGNC | ENTREZGENE_ACC |
| 3992 | ENSG00000149485 | FADS1 | fatty acid desaturase 1 [Source:HGN | ENTREZGENE_ACC |
| 400165 | ENSG00000197595 | ATP11AUN | ATP11A upstream neighbor lncRNA | ENTREZGENE_ACC |
| 401 | ENSG00000165462 | PHOX2A | paired like homeobox 2A [Source:H | ENTREZGENE_ACC |
| 401036 | ENSG00000182177 | ASB18 | ankyrin repeat and SOCS box contai | ENTREZGENE_ACC |
| 402569 | ENSG00000185467 | KPNA7 | karyopherin subunit alpha 7 [Sourc | ENTREZGENE_ACC |
| 4326 | ENSG00000198598 | MMP17 | matrix metallopeptidase 17 [Source | ENTREZGENE_ACC |
| 440350 | ENSG00000233232 | NPIPB7 | nuclear pore complex interacting p | ENTREZGENE_ACC |
| 4633 | ENSG00000111245 | MYL2 | myosin light chain 2 [Source:HGNC | ENTREZGENE_ACC |

| | | | | |
|---|---|---|---|---|
| 4669 | ENSG00000108784 | NAGLU | N-acetyl-alpha-glucosaminidase [So | ENTREZGENE_ACC |
| 4924 | ENSG00000104805 | NUCB1 | nucleobindin 1 [Source:HGNC Symb | ENTREZGENE_ACC |
| 493856 | ENSG00000145354 | CISD2 | CDGSH iron sulfur domain 2 [Source | ENTREZGENE_ACC |
| 503835 | ENSG00000258873 | DUXA | double homeobox A [Source:HGNC | ENTREZGENE_ACC |
| 50640 | ENSG00000135241 | PNPLA8 | patatin like phospholipase domain | ENTREZGENE_ACC |
| 51163 | ENSG00000138231 | DBR1 | debranching RNA lariats 1 [Source:I | ENTREZGENE_ACC |
| 51264 | ENSG00000108826 | MRPL27 | mitochondrial ribosomal protein L2 | ENTREZGENE_ACC |
| 51282 | ENSG00000171222 | SCAND1 | SCAN domain containing 1 [Source: | ENTREZGENE_ACC |
| 51380 | ENSG00000139631 | CSAD | cysteine sulfinic acid decarboxylase | ENTREZGENE_ACC |
| 5143 | ENSG00000105650 | PDE4C | phosphodiesterase 4C [Source:HGN | ENTREZGENE_ACC |
| 5143 | ENSG00000285188 | PDE4C | phosphodiesterase 4C [Source:NCBI | ENTREZGENE_ACC |
| 51433 | ENSG00000089053 | ANAPC5 | anaphase promoting complex subu | ENTREZGENE_ACC |
| 51473 | ENSG00000146038 | DCDC2 | doublecortin domain containing 2 [ | ENTREZGENE_ACC |
| 5148 | ENSG00000185527 | PDE6G | phosphodiesterase 6G [Source:HGN | ENTREZGENE_ACC |
| 51497 | ENSG00000101158 | NELFCD | negative elongation factor complex | ENTREZGENE_ACC |
| 51561 | ENSG00000110944 | IL23A | interleukin 23 subunit alpha [Sourc | ENTREZGENE_ACC |
| 5157 | ENSG00000104213 | PDGFRL | platelet derived growth factor rece | ENTREZGENE_ACC |
| 5269 | ENSG00000124570 | SERPINB6 | serpin family B member 6 [Source:H | ENTREZGENE_ACC |
| 5372 | ENSG00000100417 | PMM1 | phosphomannomutase 1 [Source:H | ENTREZGENE_ACC |
| 54363 | ENSG00000101323 | HAO1 | hydroxyacid oxidase 1 [Source:HGN | ENTREZGENE_ACC |
| 5444 | ENSG00000005421 | PON1 | paraoxonase 1 [Source:HGNC Symb | ENTREZGENE_ACC |
| 54890 | ENSG00000091542 | ALKBH5 | alkB homolog 5, RNA demethylase [ | ENTREZGENE_ACC |
| 55510 | ENSG00000080007 | DDX43 | DEAD-box helicase 43 [Source:HGN | ENTREZGENE_ACC |
| 55794 | ENSG00000182810 | DDX28 | DEAD-box helicase 28 [Source:HGN | ENTREZGENE_ACC |
| 56911 | ENSG00000156265 | MAP3K7CL | MAP3K7 C-terminal like [Source:HG | ENTREZGENE_ACC |
| 56942 | ENSG00000103121 | CMC2 | C-X9-C motif containing 2 [Source:H | ENTREZGENE_ACC |
| 57104 | ENSG00000177666 | PNPLA2 | patatin like phospholipase domain | ENTREZGENE_ACC |
| 57405 | ENSG00000152253 | SPC25 | SPC25 component of NDC80 kineto | ENTREZGENE_ACC |
| 574460 | ENSG00000207869 | MIR498 | microRNA 498 [Source:HGNC Symb | ENTREZGENE_ACC |
| 57553 | ENSG00000243156 | MICAL3 | microtubule associated monooxyge | ENTREZGENE_ACC |
| 59082 | ENSG00000255501 | CARD18 | caspase recruitment domain family | ENTREZGENE_ACC |
| 6159 | ENSG00000162244 | RPL29 | ribosomal protein L29 [Source:HGN | ENTREZGENE_ACC |
| 6184 | ENSG00000163902 | RPN1 | ribophorin I [Source:HGNC Symbol; | ENTREZGENE_ACC |
| 6227 | ENSG00000171858 | RPS21 | ribosomal protein S21 [Source:HGN | ENTREZGENE_ACC |
| 6290 | ENSG00000290741 | SAA3P | serum amyloid A3, pseudogene [So | ENTREZGENE_ACC |
| 641364 | ENSG00000250033 | SLC7A11-A | SLC7A11 antisense RNA 1 [Source:H | ENTREZGENE_ACC |
| 643224 | None | None | None | |
| 643965 | ENSG00000205116 | TMEM88B | transmembrane protein 88B [Sourc | ENTREZGENE_ACC |
| 64407 | ENSG00000150681 | RGS18 | regulator of G protein signaling 18 [ | ENTREZGENE_ACC |
| 644961 | None | None | None | |
| 645191 | ENSG00000220008 | LINGO3 | leucine rich repeat and Ig domain c | ENTREZGENE_ACC |
| 645745 | ENSG00000244020 | MT1HL1 | metallothionein 1H like 1 [Source:H | ENTREZGENE_ACC |
| 645832 | ENSG00000274529 | SEBOX | SEBOX homeobox [Source:HGNC Sy | ENTREZGENE_ACC |
| 6459 | None | None | None | |
| 647107 | ENSG00000241369 | LINC01192 | long intergenic non-protein coding | ENTREZGENE_ACC |
| 64978 | ENSG00000204316 | MRPL38 | mitochondrial ribosomal protein L3 | ENTREZGENE_ACC |
| 653162 | None | None | None | |
| 653390 | None | None | None | |
| 653583 | ENSG00000176531 | PHLDB3 | pleckstrin homology like domain fa | ENTREZGENE_ACC |
| 6649 | ENSG00000109610 | SOD3 | superoxide dismutase 3 [Source:HG | ENTREZGENE_ACC |
| 6702 | None | None | None | |
| 6722 | ENSG00000112658 | SRF | serum response factor [Source:HGN | ENTREZGENE_ACC |
| 6836 | ENSG00000148248 | SURF4 | surfeit 4 [Source:HGNC Symbol;Acc | ENTREZGENE_ACC |
| 6857 | ENSG00000067715 | SYT1 | synaptotagmin 1 [Source:HGNC Syn | ENTREZGENE_ACC |
| 6917 | ENSG00000187735 | TCEA1 | transcription elongation factor A1 [ | ENTREZGENE_ACC |
| 693229 | ENSG00000207997 | MIR644A | microRNA 644a [Source:HGNC Syml | ENTREZGENE_ACC |

| | | | | |
|---|---|---|---|---|
| 693234 | ENSG00000207575 | MIR649 | microRNA 649 [Source:HGNC Symb | ENTREZGENE_ACC |
| 7020 | ENSG00000137203 | TFAP2A | transcription factor AP-2 alpha [Sou | ENTREZGENE_ACC |
| 727897 | ENSG00000117983 | MUC5B | mucin 5B, oligomeric mucus/gel-for | ENTREZGENE_ACC |
| 7429 | ENSG00000127831 | VIL1 | villin 1 [Source:HGNC Symbol;Acc:H | ENTREZGENE_ACC |
| 7474 | ENSG00000114251 | WNT5A | Wnt family member 5A [Source:HG | ENTREZGENE_ACC |
| 7881 | ENSG00000169282 | KCNAB1 | potassium voltage-gated channel su | ENTREZGENE_ACC |
| 79170 | ENSG00000167183 | PRR15L | proline rich 15 like [Source:HGNC Sy | ENTREZGENE_ACC |
| 79575 | ENSG00000127220 | ABHD8 | abhydrolase domain containing 8 [S | ENTREZGENE_ACC |
| 79933 | ENSG00000166317 | SYNPO2L | synaptopodin 2 like [Source:HGNC S | ENTREZGENE_ACC |
| 81533 | ENSG00000129636 | ITFG1 | integrin alpha FG-GAP repeat conta | ENTREZGENE_ACC |
| 81544 | ENSG00000158555 | GDPD5 | glycerophosphodiester phosphodie | ENTREZGENE_ACC |
| 81697 | ENSG00000168131 | OR2B2 | olfactory receptor family 2 subfamil | ENTREZGENE_ACC |
| 83478 | ENSG00000138639 | ARHGAP24 | Rho GTPase activating protein 24 [S | ENTREZGENE_ACC |
| 8408 | ENSG00000177169 | ULK1 | unc-51 like autophagy activating kir | ENTREZGENE_ACC |
| 84519 | ENSG00000111644 | ACRBP | acrosin binding protein [Source:HG | ENTREZGENE_ACC |
| 84886 | ENSG00000119280 | C1orf198 | chromosome 1 open reading frame | ENTREZGENE_ACC |
| 84923 | ENSG00000133193 | FAM104A | family with sequence similarity 104 | ENTREZGENE_ACC |
| 84955 | ENSG00000120526 | NUDCD1 | NudC domain containing 1 [Source: | ENTREZGENE_ACC |
| 85376 | ENSG00000275793 | RIMBP3 | RIMS binding protein 3 [Source:HGI | ENTREZGENE_ACC |
| 8870 | ENSG00000137331 | IER3 | immediate early response 3 [Source | ENTREZGENE_ACC |
| 8871 | ENSG00000078269 | SYNJ2 | synaptojanin 2 [Source:HGNC Symb | ENTREZGENE_ACC |
| 8879 | ENSG00000166224 | SGPL1 | sphingosine-1-phosphate lyase 1 [S | ENTREZGENE_ACC |
| 90007 | ENSG00000167470 | MIDN | midnolin [Source:HGNC Symbol;Acc | ENTREZGENE_ACC |
| 9002 | ENSG00000127533 | F2RL3 | F2R like thrombin or trypsin recept | ENTREZGENE_ACC |
| 90523 | ENSG00000146147 | MLIP | muscular LMNA interacting protein | ENTREZGENE_ACC |
| 9092 | ENSG00000175467 | SART1 | spliceosome associated factor 1, re | ENTREZGENE_ACC |
| 91608 | ENSG00000270885 | RASL10B | RAS like family 10 member B [Sourc | ENTREZGENE_ACC |
| 93492 | ENSG00000132958 | TPTE2 | transmembrane phosphoinositide 3 | ENTREZGENE_ACC |
| 9779 | ENSG00000131374 | TBC1D5 | TBC1 domain family member 5 [Sou | ENTREZGENE_ACC |
| 9955 | ENSG00000153976 | HS3ST3A1 | heparan sulfate-glucosamine 3-sulf | ENTREZGENE_ACC |
